# Evolution towards higher unitary conductance in mammals makes BK channels more efficient and precise

**DOI:** 10.1101/2025.10.05.680598

**Authors:** Iain Hepburn, Tomohiko Taniguchi, Erik De Schutter

## Abstract

The large conductance calcium-activated potassium channel, known as the BK channel, play an essential role in neuronal firing and is characterized by a very large ∼250pS unitary conductance in mammals. However, this high unitary conductance is not consistent across all species with invertebrates demonstrating a much lower unitary conductance. We explored the calcium activation properties of BK channels of different unitary conductance in computational models and found that mammalian channels are more efficiently activated by calcium and produce a stronger potassium current compared to the channels of lower unitary conductance found in invertebrates. The lower unitary conductance channels display fierce competition for the available calcium, which results in low activation and weaker current. Due to these properties, mammalian BK channels are more suitable to repolarize sodium action potentials, which enables more precise spike timing, and may explain why evolution appears to have favored a trend towards very high unitary conductance in mammalian BK channels. This may be an essential component of the more advanced brain functions achieved by these species compared to invertebrates.

## Introduction

Neurons generate electrical signals via the activity of ion channels embedded in their membranes, which conduct currents selectively and may be activated by voltage changes across the membrane or by binding ligand molecules. The “Big Potassium” channel (BK channel, encoded by the Slo-1 gene^1^) has a special property in that it is activated both by a depolarizing voltage and internal calcium. Expressed extensively throughout the central nervous system, it plays an essential role in cell excitability and neurotransmitter release^2,3^, regulates neural firing^3,4^, and both excessive or insufficient activity of this channel is implicated in many neurological disorders such as epilepsy, ataxia and intellectual disability^4,5^.

BK channels consist of four identical subunits which form a pore that conducts potassium current selectively across the cell membrane at a characteristic single-channel, or unitary, conductance when the channel is activated. This unitary conductance describes the current that one single ion channel will conduct across the membrane at typical physiological conditions. The BK channel is characterised by a strikingly large unitary conductance compared to other types of ion channel^6.^

BK channels require high, micromolar levels of calcium to activate, which is achieved by close localisation to calcium-permeable channels such as the P-type calcium channel (CaP)^7,8^. The nanodomains that form around these clusters in which calcium is sufficiently high to activate the channel are small in size, of the order of 10nm^9^. The CaP channel is activated by voltage and conducts a calcium current mainly from outside the cell to inside, which is a depolarizing current. The BK potassium channel is activated both by voltage and the calcium influx from the CaP channel, and conducts a potassium current from inside the cell to outside, repolarising the cell (Fig. 1b).

**Fig. 1:**
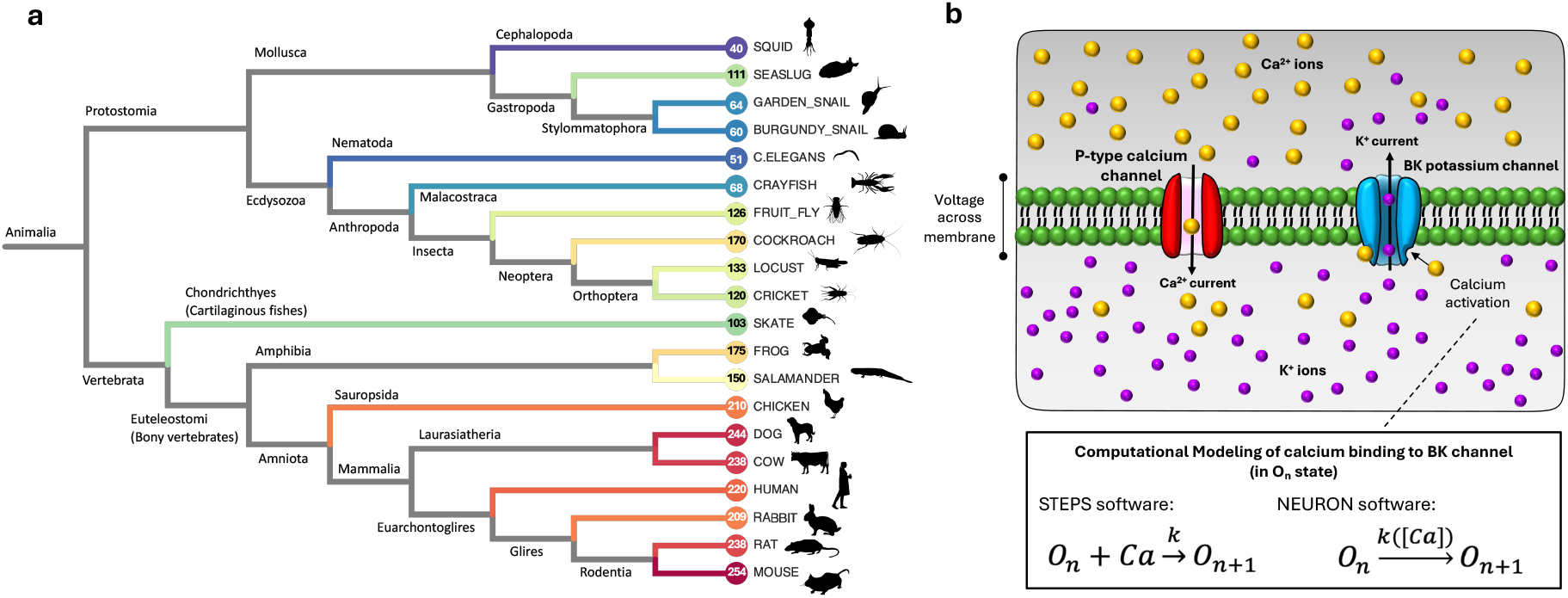
BK unitary conductance variation across animal species, and the action of BK channels and CaP channels in the membrane. **a**, A phylogenetic tree of animal species with reported BK unitary conductance in literature, as shown in Table 1. **b**, A schematic of the two ion channels described in this study. The BK channel consists of four α subunits (corresponding to n in the reaction scheme), each of which can bind a Ca^2+^ ion, leading to an increasing likelihood of being in the conducting state with greater n^25^ (Supplementary Fig. 1). The calcium activation of the BK channel is explicitly modeled molecularly in the STEPS simulator, involving a binding reaction, but in the NEURON simulator it is modeled implicitly whereby the calcium concentration modifies the activation reaction rates.

The importance of BK channels in healthy neuronal functioning is apparent by their persistence throughout millions of years of evolution (Fig. 1a and Table 1), however, BK channel unitary conductance has not been preserved throughout evolution; an interesting change has occurred. In Protostomia invertebrates, which last shared a common ancestor with vertebrates ∼560 million years ago^10^, the unitary conductance of BK channels is low, but already higher than other potassium channels^6^ . In molluscs and nematodes unitary conductance is typically observed to be less than or up to 100 picosiemens (pS) ^11-14^; in insects larger at between 100pS and 200pS^15-17^. Within vertebrates, there is a clear trend towards higher unitary conductance, always greater than 200pS in mammals and birds^18-24^. Thus, the unitary conductance of mammalian BK channels is outstandingly high when compared with other species or with any other ion channel^6^, which is due to specific adaptations of the channel pore^13^. So why has this evolutionary trend towards a higher unitary conductance occurred? To investigate, we applied two computational models: the first a stochastic molecular model with discrete ion channels, by which unitary conductance could easily be modified, in which we extensively investigated the activity of BK channels of different unitary conductances in axonal morphology. We then also used a published whole Purkinje cell model to investigate how BK channel unitary conductance affects neuronal spike timings in these cells, and discuss how the calcium buffering properties of these channels may be an important factor that has driven evolution of this channel towards higher unitary conductance in mammals.

**Table 1:** Reported BK channel unitary conductance for different animal species in the literature.

| Name | Scientific Name | BK channel unitary conductance (pS) | Average (pS) |
| --- | --- | --- | --- |
| Longfin Inshore Squid | *Doryteuthis pealeii* | 40<sup>26</sup> | 40 |
| California sea hare | *Aplysia californica* | 104<sup>13</sup>, 117<sup>27</sup> | 111 |
| Garden snail | *Cornu aspersum* (*helix aspersa*) | 64<sup>11</sup> | 64 |
| Burgundy snail | *Helix pomatia* | 60<sup>12</sup> | 60 |
| *C. elegans* | *Caenorhabditis elegans* | 51<sup>14</sup> | 51 |
| Red swamp crayfish | *Procambarus clarkii* | 68<sup>28</sup> | 68 |
| Fruit fly | *Drosophila melanogaster* | 126<sup>15</sup> | 126 |
| American cockroach | *Periplaneta americana* | 170<sup>16</sup> | 170 |
| Desert locust | *Schistocerca gregaria* | 133<sup>29</sup> | 133 |
| Two-spotted cricket | *Gryllus bimaculatus* | 120<sup>17</sup> | 120 |
| Little skate | *Leucoraja erinacea* | 103<sup>30</sup> | 103 |
| African clawed frog | *Xenopus laevis* | 175<sup>31</sup> | 175 |
| Tiger salamander | *Ambystoma tigrinum* | 150<sup>32</sup> | 150 |
| Leghorn chicken | *Gallus domesticus* | 210<sup>18</sup> | 210 |
| Dog | *Canis lupus familiaris* | 244<sup>19</sup> | 244 |
| Cattle | *Bos taurus* | 238<sup>20</sup> | 238 |
| Human | *Homo sapiens* | 220<sup>33</sup>, 229<sup>34</sup>, 210<sup>35</sup>, 221<sup>35</sup>, 220<sup>21</sup> | 220 |
| Rabbit | *Oryctolagus Cuniculus* | 230<sup>36</sup>, 200<sup>37</sup>, 180<sup>38</sup>, 226<sup>39</sup> | 209 |
| Fancy Rat | *Rattus norvegicus domestica* | 231<sup>22</sup>, 220<sup>40</sup>, 220<sup>41</sup>, 279<sup>42</sup>, 242<sup>23</sup> | 238 |
| House Mouse | *Mus musculus* | 276<sup>24</sup>, 237<sup>43</sup>, 269<sup>30</sup>, 250<sup>44</sup> | 254 |

## Results

### Animal BK channel unitary conductance

We performed a thorough literary search of reports of BK channel unitary conductance across different animal species. Table 1 shows our findings. We then constructed a phylogenetic tree of this data, as shown in Fig. 1a. What we found was a stark difference in BK unitary conductance between different phyla and classes of animals. Invertebrates, with reports from molluscs, nematodes, insects and decapods, all showed a unitary conductance below 200pS. Molluscs and nematodes ranging from 40pS up to 111pS. Insects were distinctly higher, ranging between 120pS and 170pS.

Amongst vertebrates, cartilaginous fish diverged from other vertebrates ∼460M years ago^10^ and demonstrate a low unitary conductance reported at 103pS in the Skate. Amphibians are also clearly distinguished from amniotes (which includes mammals and birds) with unitary conductance ranging between 150pS and 175pS, whereas mammals and birds all have values above 200pS, reaching up to ∼280pS for rodents. We set ourselves the task of exploring, in computational models, the influence of BK channel unitary conductance in spiking properties of neurons to try to ascertain why evolution seems to have gone hand in hand with a change in BK unitary conductance across major animal classes and phyla, with a clear trend towards higher unitary conductance in insects and vertebrates, being particularly high for mammals and birds.

### Calcium-activation of BK channels scales poorly compared to voltage-activation of CaP channels, with the worst performance in low unitary conductance channels

We took a model of a Purkinje cell^45^ originally written in NEURON^46^ and previously converted to STEPS^47^ for molecular simulation^48^. STEPS is a molecular simulator in which we have previously shown how noise in BK channels plays a key role generating dendritic calcium spike variability in Purkinje cells^49^. For simplicity, we took only the axonal section of the model and ran in a cylinder geometry of 1µm diameter and 10µm length. The model included (amongst other ion channels^50^) voltage-activated P-type calcium channels (CaP channels) and calcium-activated BK channels, along with calcium buffers and pumps. Since STEPS is a discrete molecular simulator (i.e. based on integer number of molecules and not floating-point concentrations), conversion to a STEPS model requires defining a unitary (single-channel) conductance. When total BK conductance in membrane is conserved, varying the BK channel conductance varies the number of BK channels present on the membrane, i.e. a lower unitary conductance gives a higher number of channels in the membrane. For example in this model, a 50pS unitary conductance resulted in 37,356 BK channels in the membrane but only 7,471 channels when unitary conductance was 250pS, both for a total BK maximal membrane conductance of 1.87µS. For our first simulations, we ran for a short biological time, 5ms, in which the first spontaneous spike appears, and show results from these two channels in Fig. 2.

**Fig. 2:**
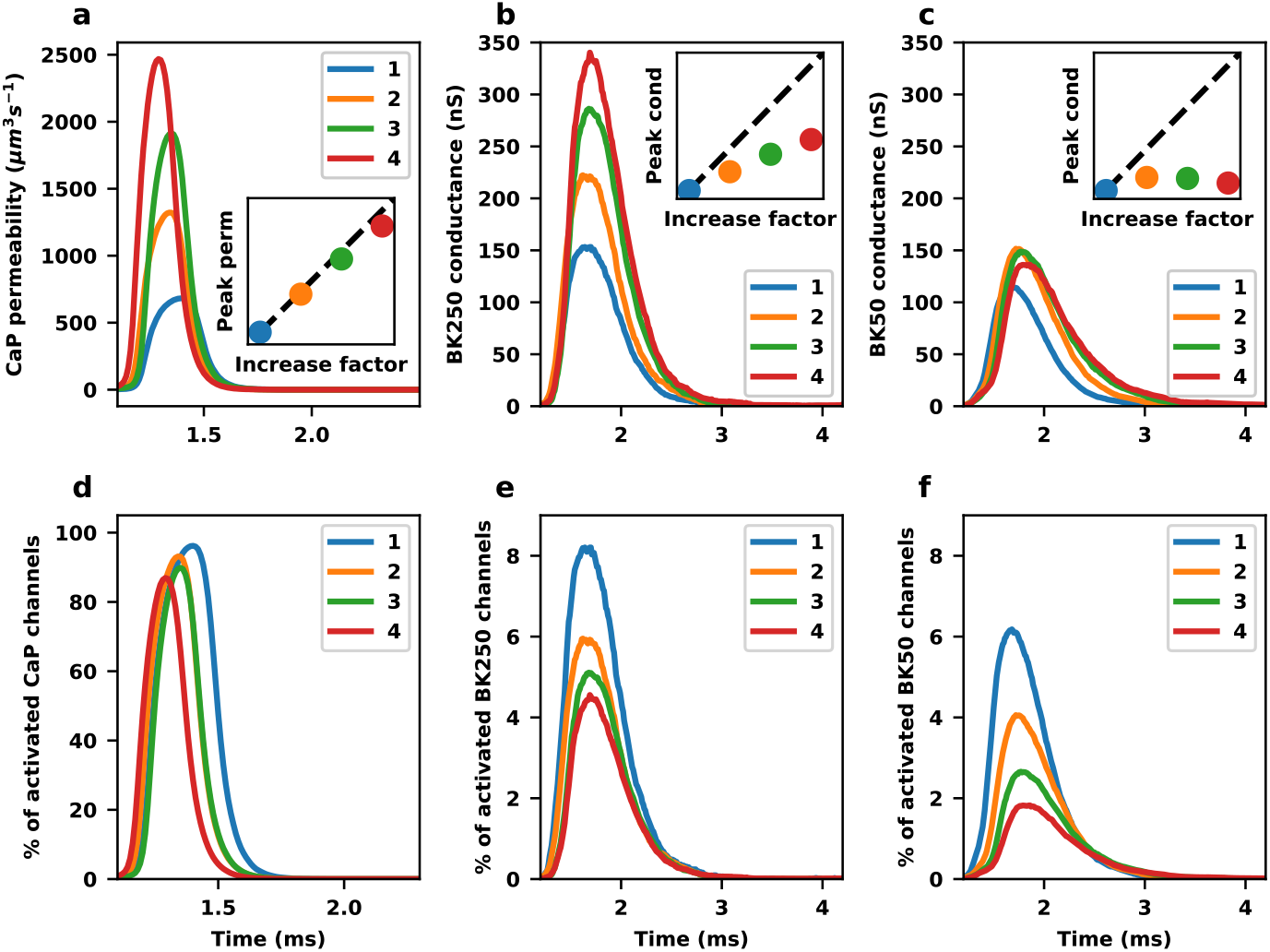
Activation of CaP and BK channels in a discrete molecular computational model. **a**, Total CaP membrane permeability (single-channel permeability multiplied by the number of open channels) with maximal CaP permeability (the membrane permeability if all channels are open) increased by factors of 2 (orange), 3 (green), 4 (red) and compared to the original model (factor 1, blue). The inset shows the peak permeability for the 4 different factors normalised to the peak permeability in the factor=1 case, and compared to the ‘ideal’ case where peak permeability increases proportionally to the increase factor. **b**, Total BK membrane conductance (single-channel conductance multiplied by the number of open channels) in a model with BK unitary conductance of 250pS and maximal BK conductance (the membrane conductance if all channels are open) increase factors of 1, 2, 3, 4 (inset is the same as for **a**). **c**, Total BK conductance in model with BK unitary conductance of 50pS and maximal BK conductance increase factors of 1, 2, 3, 4 (inset is the same as for **a**). **d**, in the model where CaP maximal permeability is systematically increased, the percentage of activated CaP channels that lead to the permeability shown in **a. e**, In the 250pS unitary conductance BK channel model, the percentage of activated BK channels that lead to the conductance shown in **b. f**, In the 50pS unitary conductance BK channel model, the percentage of activated BK channels that lead to the conductance shown in **c**. All simulations were run with STEPS.

We first increased the maximal CaP permeability by factors of 2, 3, and 4 (by increasing the number of channels by this factor) and compared to the original model (factor = 1). As expected, we observed a near linear increase in peak CaP permeability (Fig. 2a), demonstrated by a high percentage of channels activated in all cases (Fig. 2d). However, when we also increased the BK maximal conductance by the same factors, there was a stark difference to the CaP behavior. When BK unitary conductance was 250pS, an increase in maximal BK conductance did increase the peak current, but sublinearly (Fig. 2b). This shows in a significantly lower activation of BK channels from ∼8% when factor was 1 to only ∼4% with an increase factor of 4 (Fig. 2e). When BK unitary conductance was 50pS the difference from the CaP channel was even starker, as can be seen in Fig. 2c. Here, with any increase in total BK conductance there was no resulting increase in peak BK conductance and in fact even a slight drop. This shows in a strong fall in percentage of channels activated from ∼6% of all channels activated with factor of 1 to under 2% with increase factor of 4 (Fig. 2f).

The only difference between the simulations comparing Fig. 2b to 2c or 2e to 2f is the number of BK channels present on the membrane surface and their unitary conductance, and the most important difference between BK channels and CaP channels (which show much stronger activation compared to the BK channels), is the calcium-activation of BK channels. Therefore, we next recorded calcium binding to channels and submembrane calcium to try to explain the effects observed.

### Lower activation of BK channels with low unitary conductance is caused by their buffering of calcium, effectively causing competition between channels

Since a 4-fold increase in maximal BK conductance also means a 4-fold increase in the number of BK channels, the poorer activation observed in BK channels (Fig. 2e,f) compared to CaP channels (Fig. 2d) at higher increase factors hints at a significant buffering effect of the BK channels, i.e. channels are removing calcium from the system and this is reducing the activation of more channels. It can be expected that the buffering effect depends on the unitary conductance, because varying the unitary conductance changes the number of channels available to bind calcium. This is of course not a concern for CaP channels, which are only activated by voltage.

To explore this buffering effect of calcium by BK channels, and how the strength of this effect may depend on the unitary conductance, we ran our model for 5 different 100ms simulations, varying only the BK channel unitary conductance between simulations (whilst maintaining the same maximal membrane BK conductance). The simulations generated spontaneous spikes, aligned on an initial early spike (inset of Fig. 3a) then generating later spikes with different timings (Fig. 3a) due to different firing properties induced by the BK unitary conductance change and stochastic effects. We then measured total BK conductance on the membrane (Fig. 3b and Fig. 3c show maximum BK membrane conductance over 100ms), calcium ions bound to BK channels (Fig. 3d) and the submembrane calcium concentration (Fig. 3e,f). The simulations confirmed that the lower unitary conductance channels were less able to activate and generate BK current. Fig. 3b,c shows that a unitary conductance of 250pS (approximately the unitary conductance of mammals, Fig. 1b) results in peak BK conductance of approximately 155nS compared to only 120nS with a unitary conductance of 50pS (approximately the unitary conductance of non-insect invertebrates, Fig. 1b). Examining the calcium profiles elucidates why. As shown in Fig. 3d and as can be expected, more calcium ions bind to BK channels when their unitary conductance is low (and therefore the number of channels is high), which reduces the peak submembrane calcium concentration (Fig. 3e), meaning that less calcium is available to further activate channels. In contrast, when unitary conductance is high a much lower number of calcium ions binds to channels to activate the BK current (Fig. 3d), resulting in a higher submembrane calcium concentration (Fig. 3e) and therefore stronger activation and a stronger BK current (Fig. 3b). In addition, slower unbinding of calcium from the low unitary conductance channels (Fig. 3d) results in a wider submembrane calcium profile (Fig. 3e,f). In summary, high unitary conductance channels give higher and narrower calcium peaks (Fig. 3f) leading to a higher BK membrane conductance (Fig. 3b) compared to low unitary conductance channels. This is a result of a lower buffering effect on calcium (Fig. 3d), effectively a more efficient use of calcium. These findings are described schematically in Fig. 3g.

**Fig. 3:**
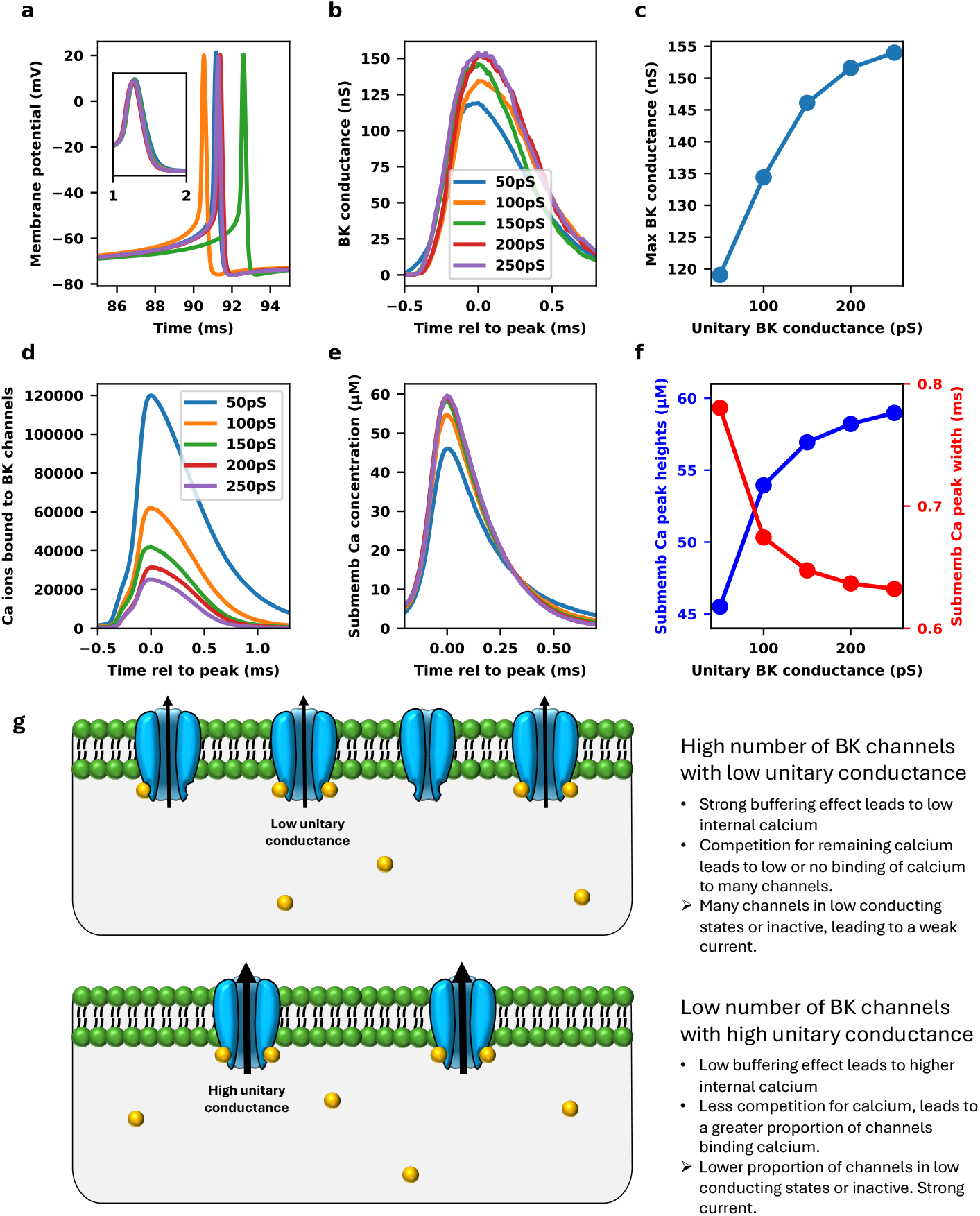
Calcium activation of BK channels of different unitary conductance. **a**, Spontaneous spiking in the simulations, with an initial aligned spike (inset) and later spike timings depending on unitary conductance and stochastic effects (each different trace shows a different unitary conductance). **b**, BK membrane conductance aligned on the peak conductance over the 100ms simulation for the different unitary conductance channels tested ranging from 50pS (blue) to 250pS (purple). **c**, The maximum BK membrane conductance over the 100ms simulation plotted vs unitary conductance ranging from 50pS to 250pS. **d**, The number of calcium ions bound to BK channels aligned on maximum over the 100ms simulation, of the different BK unitary conductances. **e**, Submembrane calcium concentration for the simulations aligned on peak over the 100ms simulations. **f**, Submembrane calcium peak height (blue) and width (red) plotted vs BK channel unitary conductance over the 50pS-250pS range. The width is calculated at a relative height of 10% of peak amplitude. All simulations in **a**-**f** were run with STEPS. **g**, A schematic diagram with accompanying text explaining the effects observed.

### Modified BK channels that do not bind calcium ions behave similarly to voltage-activated channels

To confirm that the buffering of calcium is the main cause of the differences between the behavior of BK channels compared to voltage-activated CaP channels, we created simulations whereby BK channels were still activated by calcium, as normal, but did not remove a calcium ion from the compartment during the reaction. In effect, even though the channels are activated by calcium they do not act as a buffer. Supplementary Fig. 2 shows results from this model for the same simulations as shown in Fig. 3, demonstrating clearly that there is no buffering effect of calcium from this BK channel, i.e. submembrane calcium concentration did not depend on unitary conductance.

We compared this to the normal situation where BK channels do buffer calcium, and again varied the unitary conductance. In addition, for comparison we included simulations where the CaP channel unitary permeability was varied (and so too, correspondingly, the number of CaP channels). For each set of simulations, we systematically increased the maximal conductance on the membrane. Fig. 4a left panel shows the behavior of the standard model (similar to Fig. 3b), then to the right we increased maximal BK conductance (G_BK) up to a factor of 5 (i.e. we increased the number of channels by a factor of 5). As expected, the effect of the increase in total conductance depended on the unitary conductance (as previously shown in Fig. 2b,c). A 5-fold increase resulted in peak BK membrane conductance from 250pS unitary conductance channels going from ∼150nS to nearly 400nS, but for 50pS channels barely resulted in any increase at all (Fig. 4a).

**Fig. 4:**
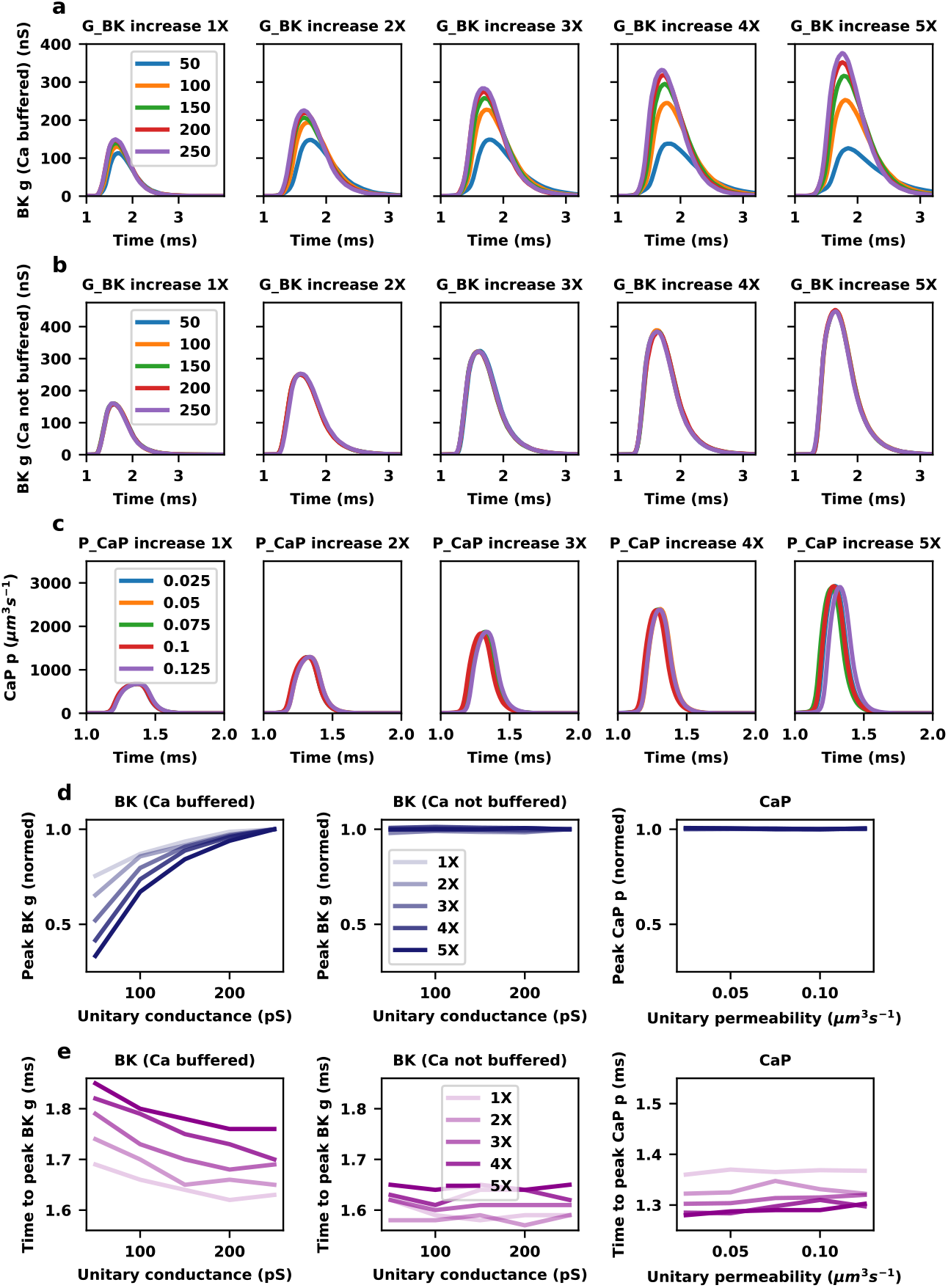
Comparison between BK channels that bind calcium, BK channels that do not bind calcium, and voltage-activated CaP channels. **a**, Activation of BK channels that bind calcium. To the right the maximal BK conductance is increased up to a factor of 5, and for each set of simulations BK unitary conductance was varied from 50pS (blue) to 250pS (purple). **b**, Activation of BK channels that do not bind calcium. To the right the maximal BK conductance is increased up to a factor of 5, and for each set of simulations BK unitary conductance varied from 50pS (blue) to 250pS (purple). **c**, Activation of CaP channels. To the right the maximal CaP permeability is increased up to a factor of 5, and for each set of simulations CaP unitary permeability was varied from 0.025µm^3^s^-1^ (blue) to 0.125 µm^3^s^-1^ (purple). **d**, The peak membrane conductance or permeability (normalized to maximum) shown separately for the different increase factors from 1 (lightest blue) to 5 (darkest blue) and plotted vs unitary conductance or permeability over the ranges tested, for the 3 different types of channels. **e**, The time to peak membrane conductance shown separately for the different increase factors from 1 (lightest purple) to 5 (darkest purple) and plotted vs unitary conductance or permeability over the ranges tested, for the 3 different types of channels. All simulations were run with STEPS.

In simulations where BK channels did not buffer calcium but were still activated by it (similar to the behavior in simulators such as NEURON, Fig. 1b), results were very different. In all simulations, peak BK membrane conductance increased more strongly with an increase in maximal BK conductance compared to channels that bound calcium from ∼150nS to ∼450nS (Fig. 4b) and there was no significant effect of unitary conductance. The behavior of BK channels in simulations where they do not buffer calcium more closely resembles voltage-activated channels, such as CaP (Fig. 4c, Fig. 2a). This is demonstrated quite clearly when comparing the peak membrane conductance (Fig. 4d) and time to peak (Fig. 4e) between the 3 different channels, whereby the BK channels that don’t buffer calcium and the CaP channels have no dependence on the unitary conductance or permeability. For only the BK channel that buffer calcium the activation depends on unitary conductance, and for all cases the 50pS channel had the weakest and slowest response, performing worse as maximal conductance was increased. Similar results were observed for BK channel activation with different clamped calcium concentrations (Supplementary Fig. 3).

This all serves to demonstrate that the activation of BK channels is limited by the buffering effect of binding calcium, and this effect is stronger for lower unitary conductance channels. It is not simply because there are more channels to activate with lower unitary conductance because the simulations in Fig. 4b (where there is no effect of unitary conductance) contain the exact same number of channels as the corresponding simulations shown in Fig. 4a (where there is). It is the buffering effect reducing the submembrane calcium (Fig. 3f,g) that is important.

### Calcium buffering by BK channels affects neuronal firing properties, with low unitary conductance channels demonstrating less reliable spike timing in a Purkinje cell model

BK channels play a crucial role in neuronal spike timing^2-4^, so the buffering effect of calcium on BK channels at different unitary conductances described in this study may have a significant influence on the firing properties of neurons. BK channels are widely expressed throughout the brain, but to investigate just one role in an available computational model, we used a published Purkinje cell model in NEURON^50^. Since NEURON does not explicitly model the binding of calcium in BK activation (Fig. 1b) we added a buffering reaction to the model to mimic this effect, and by varying the concentration of this buffer we were able to effectively model different BK unitary conductances (Supplementary Fig. 4). The model contains random background synaptic noise and generates spikes spontaneously^50^. By injecting current into the soma the firing rate can be modified, and we injected current over a range of values in two different models with effective unitary BK conductances of 50pS and 250pS. Fig. 5a shows raster plots of 500 trials demonstrating mean inter-spike intervals (ISIs) of 50ms with the 50pS BK channel (left panel) and with the 250pS BK channel (right panel). We measured the ISIs and investigated two different scenarios where ISI mean was equal or injected current was equal. Under all scenarios, the coefficient of variation (CV) of the spike ISIs in the model with 50pS BK channels was greater than with 250pS channels. Fig. 5b shows an example comparison aligned on mean ISI of 50ms. The CV in the 250pS model is 0.39, whereas it is far higher at 0.66 in the 50pS model. CV was always higher in the 50pS model compared to the 250pS model over a range of mean ISIs from 20ms to 50ms (Fig. 5d). This was not some artifact of the different current injections required to get matching mean ISIs, because CV remained higher in the 50pS model even with equal current injections over a range of -0.2nA to 0.2nA (Fig. 5e). At 0.2nA in the 50pS model the cell starts to enter burst-pause behavior, which is why the CV increases very sharply towards 1 at this point and beyond. The cell does not exhibit burst-pause behavior before this point, the higher CVs in the 50pS from lower current injections is with regular spiking (e.g. Fig. 5c).

**Fig. 5:**
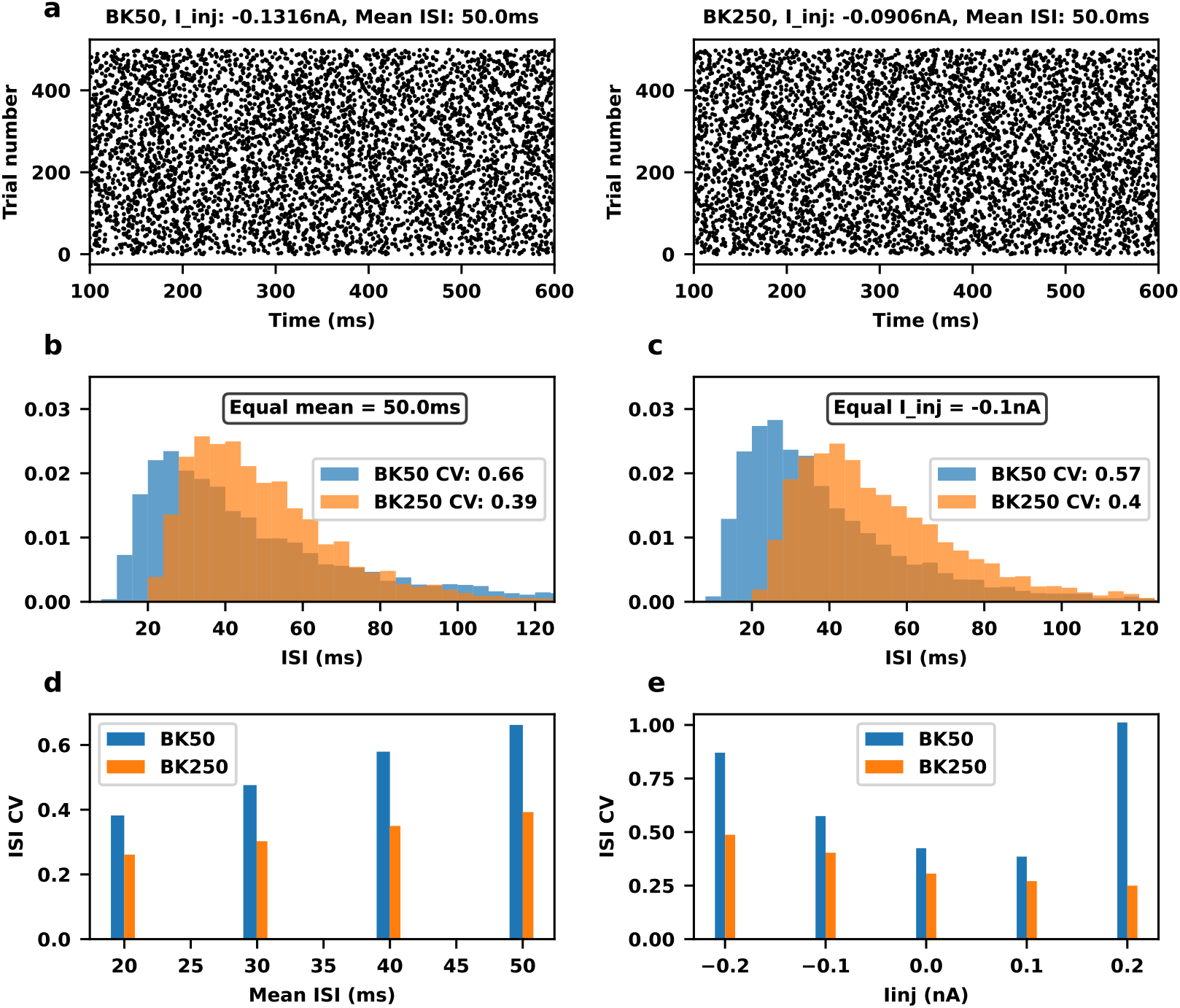
Purkinje cell activity in NEURON simulations with calcium buffering added to model BK unitary conductance of 50pS and 250pS. **a**, Raster plot of spike times in the model with effectively a 50pS (left panel) and 250pS (right panel) BK unitary conductance for 500 trials, both with a mean ISI of 50ms. **b**, normalized ISIs over all 500 trials with current injected to give a mean ISI of 50ms for both the 50pS (blue) and 250pS (orange) model. **c**, normalized ISIs over all 500 trials with equal current injection of -0.1nA in both the 50pS (blue) and 250pS (orange) model. **d**, ISI CV for different current injections to give a range of mean ISIs from 20ms to 50ms for both the 50pS (blue) and 250pS (orange) model. **e**, ISI CV for equal current over a range of -0.2nA to 0.2nA for both the 50pS (blue) and 250pS (orange) model.

The buffering effect of calcium and corresponding reduction in submembrane calcium appears to increase variance in Purkinje cell firing leading to less reliable spike times, and the effect is stronger with lower unitary conductances. This is consistent across a range of mean firing rates and current injections.

## Discussion

The BK channel is characterized by a high single-channel conductance and activation both by voltage and calcium. It consists of a tetramer of pore-forming *α* subunits, which are encoded by the Slo gene family, and the specific structure of the channel determines its conductance. Besides the 10-20 times higher single channel conductance, it has several unique properties compared to Kv channels and ion channels in general. Structural analysis shows its voltage-sensor domain to interact with the pore domain within the same subunit, while in Kv channels the voltage-sensor latches to neighboring subunits^13^. Calcium binding causes conformational changes to the Ca^2+^ sensor that affect both pore and voltage-sensor domains^51^. While the K^+^ selectivity filter is similar to that of Kv channels, the pore expands greatly below it and has an electronegative surface that attracts K^+^ ions, explaining the much larger conductance of a single BK channel^13^. It is, therefore, not surprising that analysis of the amino acid sequence of BK channels suggests that they separated early from Kv channels in evolution^52^. Moreover, in mammals BK channels evolved structurally by obtaining β4 subunits^53^ that form a tetrameric extracellular crown over the pore ^54^.

While the structural basis of the large BK conductance is known^13^ its evolutionary origin has not been described before. In this study we first collected reports of BK unitary conductance from different animal species and found that in nature these channels display a wide variation across different animal groups. Despite the low unitary conductances we include in our analysis from molluscs and other non-insect invertebrates these channels are considered BK channels due to their pharmacology and gating properties such as activation both by internal calcium and membrane depolarization^13,28,55,56^, and are distinct from the lower unitary conductance SK and IK channels that are activated only by internal calcium^57-59^. The Aplysia BK channel, for example, has a common sequence identity of 59% to the human BK channel^13^ and exhibits all the same pharmacological properties as the human BK channel yet with a lower conductance of ∼110pS compared to ∼220pS for humans (Table 1). Similarly, the coding region of the drosophilia slo-1 gene is ∼60% identical to the mouse slo-1 gene^60^ yet unitary conductance of the drosophilia BK channel is approximately half that of the mouse (Table 1). Other invertebrates also have low BK unitary conductances such as 51pS for C. elegans, whereas all unitary conductances reported from mammals and birds are over 200pS. So it seems the main evolution in this channel in vertebrates after diverging from invertebrates hundreds of millions of years ago has been towards high unitary conductance, whilst maintaining the same pharmacological properties.

We explored, computationally, the effect of calcium binding to BK channels and the differences in activation that this effect causes in these channels compared to voltage-activated channels. More calcium binds to BK channels the more channels there are, which essentially means BK channels act as a calcium buffer. This buffering effect can be expected to be on a different magnitude for different species with less buffering in species with high unitary conductance channels and more buffering in species with low unitary conductance channels. Essentially, lower unitary conductance channels require more calcium to activate compared to higher unitary conductance channels (Supplementary Fig. 3) and this affects the ability of the cell to generate BK current because there is more competition with lower unitary conductance. This may have profound effects on the ability for neurons to generate signals with low unitary conductance calcium-activated channels. We demonstrated one case in this study where the calcium buffering effect resulted in less reliable spike timing with higher variation in a Purkinje cell model with 50pS unitary conductance BK channels, while the model with 250pS channels displayed a much lower variation.

This effect of single channel conductance on spike timing (Fig. 5), may be the evolutionary pressure that increased unit conductance. Many invertebrates, like *C. elegans*, lack Na^+^ channels and do not express classic fast action potentials^61^, some neurons in *C. elegans*, however, fire much slower calcium spikes^62^. It is noticeable that BK channels, which have a low single channel conductance of about 51pS (Table 1), play only a minor role in repolarizing these calcium spikes compared to Kv channels (Figure 3H of Liu et al. 2018^62^). This fits with our prediction that low unit conductance BK channels cannot generate the large currents needed for effective hyperpolarization (Fig. 2). Therefore, the initial evolutionary pressure towards higher single channel conductance may have been to make BK channels more useful to repolarize calcium spikes. Moreover, because of the large difference in time scale, spike half-width of 18 ± 0.8ms for *C. elegans* AWA neurons^62^ versus 0.15 ± 0.01ms for rat Purkinje neurons^63^, the buffering effect of BK channels on the Ca^2+^ concentration is less relevant for calcium spike repolarization. *Aplysia*, which has mixed Na^+^-Ca^2+^ action potentials^64^ with a half-width of more than 10 ms^65^ expresses BK channels with a higher unit conductance of about 111pS (Table 1).

Finally, invertebrate species that have neurons that can fire very fast action potentials, like the H1 neuron in the fly visual system that can achieve rates of 200 Hz^66^ have even higher unitary conductance values of about 126pS (Table 1). In summary, in invertebrates there is a clear correlation between the transition from slow Ca^2+^ spiking to fast Na^+^ spiking and the BK single channel conductance.

Since BK channels are so widely distributed in the brain (and indeed within other cells in the body^2^) the effects we describe here have a significant impact on brain function. We have demonstrated that BK unitary conductance has a significant effect on neuronal spike timing, with more precise timing achieved at higher unitary conductances. Some insect, bird and mammalian functions require sub-millisecond precision^67-70^, and the increase in the BK unitary conductance may have played an important role in achieving such high precision. Accurate spike timing is quite important for Purkinje cell function^71,72^, with decreased accuracy of spike timing in mouse disease models correlating with motor performance deficits^73^ (whilst absence of BK channels causes cerebellar ataxia^74^). A reduced buffering effect and therefore more efficient use of calcium may help to explain why evolution has trended towards a higher unitary conductance of BK channels in vertebrates, and this more efficient use of calcium with lower buffering appears to have gone hand in hand with brain evolution in these species.

## Methods

### BK unitary conductance and the phylogenetic tree

Measurements of unitary BK channel conductance may vary depending on asymmetric or symmetric potassium conditions^75^, and of course experimental conditions vary across the literature. For this reason, we selected reports in symmetric conditions in the range of 100uM to 180uM potassium and generally rejected asymmetric reports. The report for C. elegans shows one example of this: unitary conductance was reported as 37.9pS in asymmetrical 15mMK_out_:173mMK_in_ saline and 51.0pS in symmetrical 173mMK_out_:173mMK_in_ saline^14^, and so we took the 51pS value. Where possible, we took data from neuronal cells but due to the low number of reports for some classes of animals this wasn’t always possible. Where there was more than one report per species (Table 1), we took the average for the phylogenetic tree figure (Fig. 1a).

The phylogenetic tree was generated with Toytree^76^, with silhouette images of animals from PhyloPic^77^. OneZoom^10^ and other general web searches were used to find the common ancestors and to create the branch structure.

### Computer simulations

We based our computer simulations on a published model by Zang et al^50^. To investigate unitary conductance properties of BK channels in axonal morphology we used STEPS^47^, which is a stochastic molecular simulator that is also able to simulate membrane voltage and was recently extended to simulate vesicles and lipid rafts in a parallel MPI solution^78^. We ran all STEPS simulations in a mesh with a cylindrical boundary, of length 10µm and diameter 1µm, comprised of 13617 tetrahedrons, using only the axon part of the Zang et al model. In order to emphasize the BK current contribution to action potential depolarization we reduced the KV3 conductance and simplified its calcium component by removing SK2 and slow BK channels. We updated the BK channel with more recent gating kinetics^25^ (the Zang model uses a previous study by the same author^79^). Table 2 lists all the ion channels in the model and their conductances. This is the base model corresponding to e.g. ‘increase factor 1’ in Fig. 2 and 1X in Fig. 4, but maximal permeability of CaP and maximal conductance of BK were altered, as explained in Results. Like the original NEURON model, the STEPS model also contained a calcium pump, and parvalbumin and calbindin calcium buffers. Since the axon part of this model contains a relatively high density of channels, in many cases only one run of the model was necessary with no significant noise effects observed in the stochastic simulations. Therefore, all individual traces shown in Fig. 2, Fig. 3 and Fig. 4a,b,c are from one run of the model and not averaged. The plots shown in Fig. 4d and e are the exception and averaged from 8 runs of the model with different seeds for smoother traces.

| Ion channel | Maximal conductance or permeability | Unitary conductance or permeability | # ion channels |
| --- | --- | --- | --- |
| Kv3 | 5.76 S/cm<sup>2</sup> | 16 pS | 112068 |
| NaP | 0.0023 S/cm<sup>2</sup> | 0.37 pS | 1935 |
| Rsg (resurgent sodium) | 0.56 S/cm<sup>2</sup> | 0.764 pS | 228178 |
| CaP | 0.00228 cm/s | 0.025 μm<sup>3</sup>/s | 28390 |
| CaT | 0.000128 cm/s | 0.0000165 μm<sup>3</sup>/s | 2414915 |
| BK | 6 S/cm<sup>2</sup> | 50 – 250 pS | 37356 - 7471 |
| Ih | 0.000108 S/cm<sup>2</sup> | 0.00097 pS | 34660 |

To investigate BK unitary conductance in a Purkinje cell model we used NEURON^46^ and added calcium buffers to the original Zang et al^50^ model to mimic the calcium buffering effect of BK channels. This involved adding a buffering reaction with the same kinetics of BK channel activation, and by varying the concentration of this buffer we were able to effectively control BK unitary conductance. In total 10 buffers were added representing the 10 BK channel states (Supplementary Fig. 1), and the corresponding 26 reactions were also added to the submembrane section following the usual kinetics of BK channel activation, e.g. for the C1 to C2 calcium binding reaction:

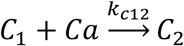

The submembrane shell width in the axon initial segment was chosen as 10nm to replicate the size of the calcium nanodomains surrounding BK channels^9^. Supplementary Fig. 4 shows the behavior of the NEURON model for 500ms runs with varying effective BK unitary conductances. As expected, the lowest unitary conductance tested (50pS) shows the strongest buffering effect with the lowest BK activation (Supplementary Fig. 4 a,b) and lowest recorded submembrane calcium concentration (Supplementary Fig. 4 c,d) .

We used Matlab code to process the spike trains shown in Fig. 5^80^. The STEPS simulations were run on a Macbook Pro with Apple M3 Max chip, and the NEURON simulations were run on the Deigo cluster at the Okinawa Institute of Science and Technology on 64-core AMD Epyc 7702 2.0GHz processors.

## Author Contributions

EDS led the study, and IH first noticed the calcium buffering effect of BK channels as the basis for this study. IH researched the phylogenetic tree. IH and TT wrote the computer code used in this study and IH ran all computer simulations. IH and EDS wrote the manuscript, and all authors read and approved of the final version.

## Acknowledgements

This work was funded by the Okinawa Institute of Science and Technology (OIST). We are grateful to the Scientific Computing and Data Analysis section of Core Facilities at OIST for providing resources that were used in this study.

