## Supplementary Material for "Evolution towards higher unitary conductance in mammals makes BK channels more efficient and precise"

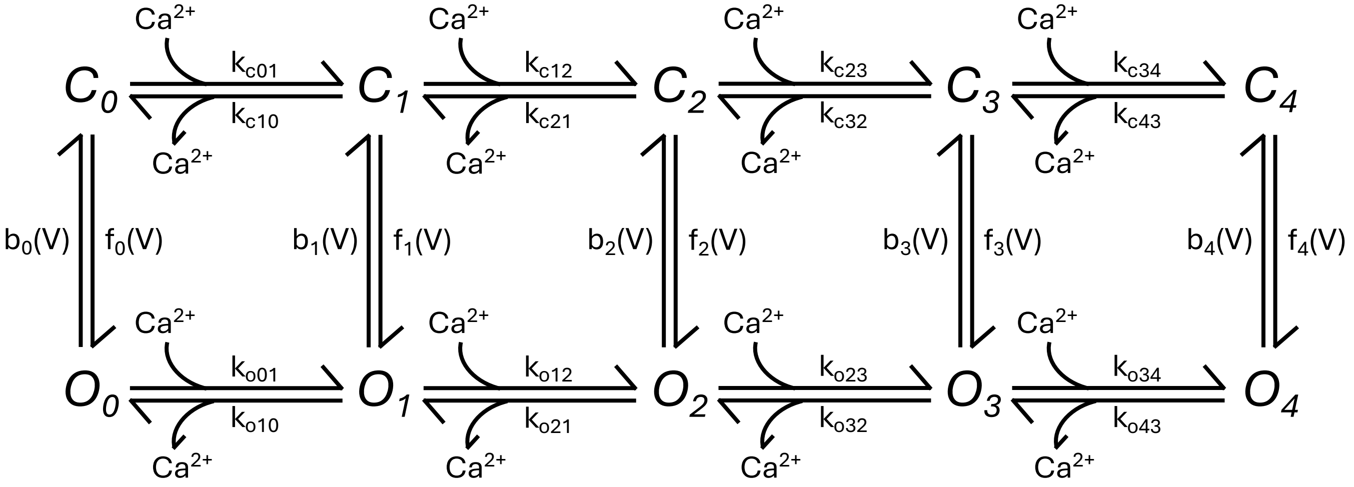


$Qt = 2^{(T-25)/10}$

$K_{c01}=K_{o01} = Qt \times 4 \times 10^9\ M^{-1}s^{-1}$
$K_{c12}=K_{o12} = Qt \times 3 \times 10^9\ M^{-1}s^{-1}$
$K_{c23}=K_{o23} = Qt \times 2 \times 10^9\ M^{-1}s^{-1}$
$K_{c34}=K_{o34} = Qt \times 1 \times 10^9\ M^{-1}s^{-1}$

$K_{c10} = Qt \times 11917 \times 1 \times 10^9\ s^{-1}$
$K_{c21} = Qt \times 11917 \times 2 \times 10^9\ s^{-1}$
$K_{c32} = Qt \times 11917 \times 3 \times 10^9\ s^{-1}$
$K_{c43} = Qt \times 11917 \times 4 \times 10^9\ s^{-1}$

$K_{o10} = Qt \times 1065 \times 1 \times 10^9\ s^{-1}$
$K_{o21} = Qt \times 1065 \times 2 \times 10^9\ s^{-1}$
$K_{o32} = Qt \times 1065 \times 3 \times 10^9\ s^{-1}$
$K_{o43} = Qt \times 1065 \times 4 \times 10^9\ s^{-1}$

$f_0(V) = Qt \times 5.5e^{0.73FV/RT}$ s-1
$f_1(V) = Qt \times 8e^{0.73FV/RT}$ s-1
$f_2(V) = Qt \times 2e^{0.73FV/RT}$ s-1
$f_3(V) = Qt \times 884e^{0.73FV/RT}$ s-1
$f_4(V) = Qt \times 900e^{0.73FV/RT}$ s-1

$b_0(V) = Qt \times 8669e^{-0.58FV/RT}$ s-1
$b_1(V) = Qt \times 1127e^{-0.58FV/RT}$ s-1
$b_2(V) = Qt \times 25.2e^{-0.58FV/RT}$ s-1
$b_3(V) = Qt \times 1013e^{-0.58FV/RT}$ s-1
$b_4(V) = Qt \times 125.7e^{-0.58FV/RT}$ s-1

*Supplementary Fig. 1: The activation scheme of the BK channel and gating kinetics.*

*The BK channel consists of 4 subunits, each of which can bind one calcium ion, and can either be closed (non-conducting) or open (conducting). There are 5 closed states (denoted by C) and 5 open states (denoted by O), each corresponding to the number of subunits with calcium bound (e.g., the C2 and O2 states both have calcium ions bound to 2 of the 4 subunits). As more calcium ions are bound, the forward voltage-activation reaction becomes faster whereas the backward reaction becomes slower- as can be seen in the kinetics- meaning the channel is more likely to be in the open state the more calcium ions it has bound.*

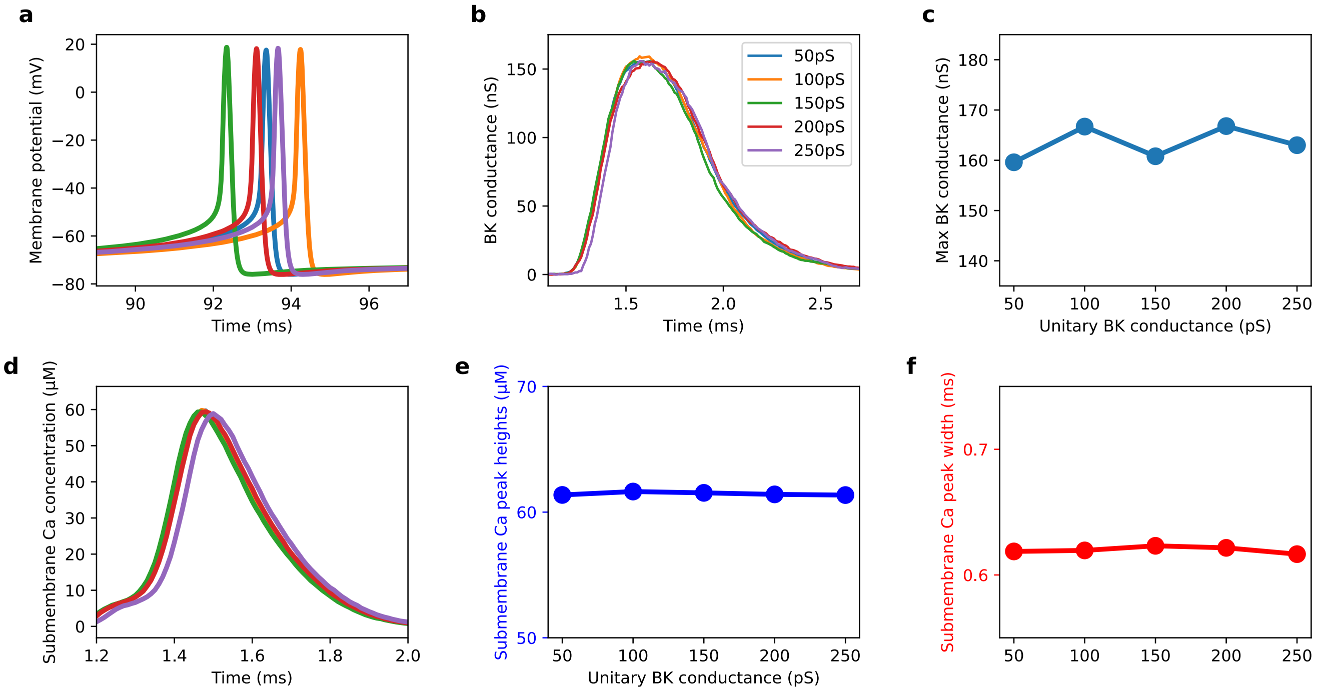


*Supplementary Fig. 2: BK current for different unitary conductances in a model where the channel does not remove calcium during activation.*

*This is the same model as shown in Fig. 3, but where the activation by calcium does not remove any calcium from the submembrane region.* ***a,*** *Spontaneous spiking for the different unitary conductances (unitary conductance key is shown in* ***b****).* ***b,*** *BK membrane conductance on the first, aligned spike.* ***c,*** *Peak BK membrane conductance over 100ms of simulation.* ***d,*** *Submembrane calcium, showing a difference to Fig. 3e in that the lower unitary conductance channels do not reduce calcium concentration in this model.* ***e,*** *Maximum submembrane calcium peak heights over the 100ms simulations.* ***f,*** *Maximum submembrane calcium peak widths over the 100ms simulations.*

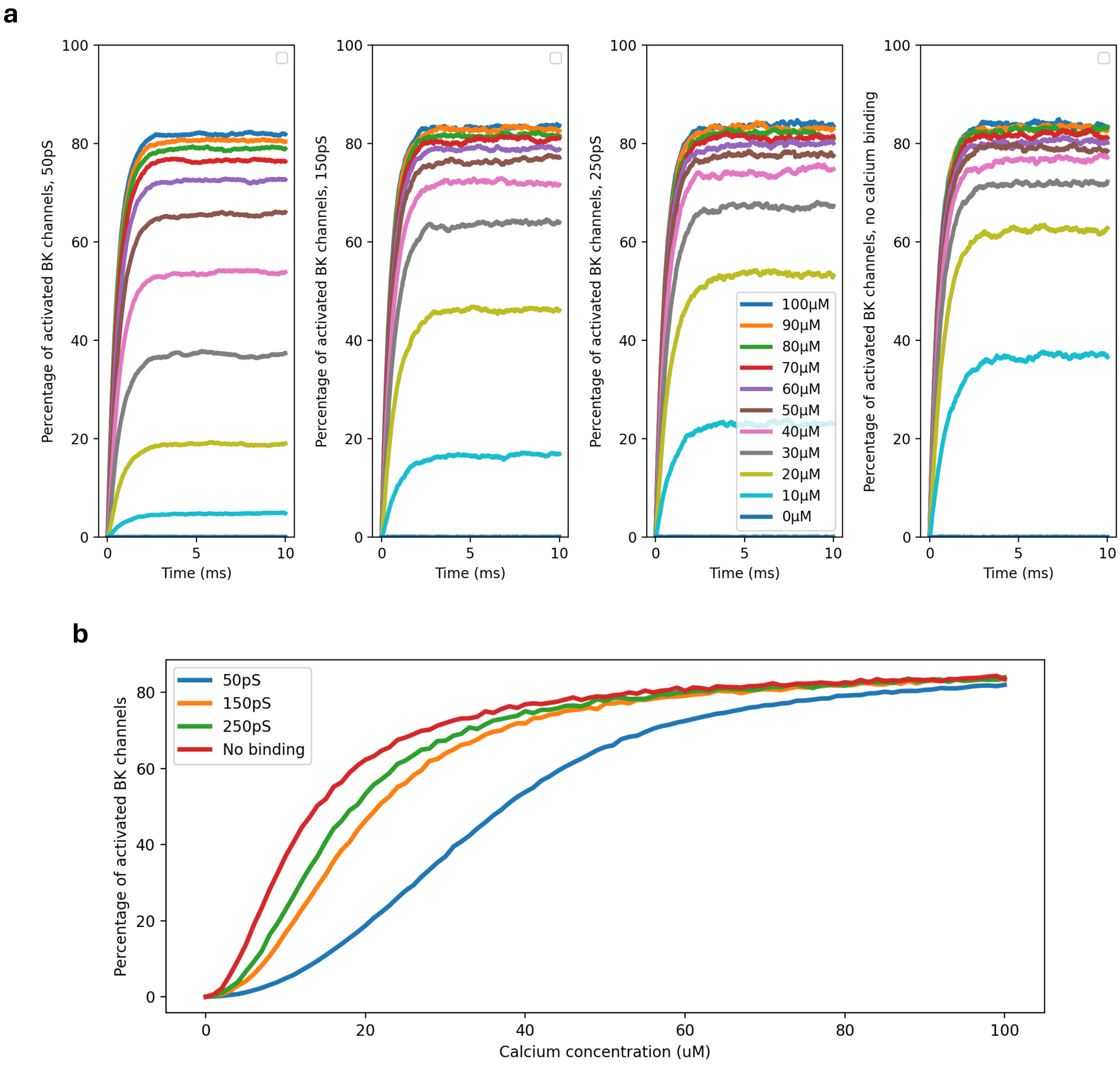


*Supplementary Fig. 3: Activation of BK channels of 50pS, 150pS and 250pS unitary conductance, and of a model BK channel that does not buffer calcium.*

*In this model that only includes a BK channel, voltage is clamped at 0mV and calcium is injected at the different levels shown, and the BK channel activates.*
***a,*** *The activation of BK channels of, from left to right, 50pS unitary conductance, 150pS unitary conductance, 250pS unitary conductance, and also 250pS but in a model whereby the BK channels does not buffer calcium.* ***b,*** *A plot summarizing the results from* ***a****, showing percentage of active BK channels vs initial calcium concentration after 10ms for the four different types of model BK channels shown.*

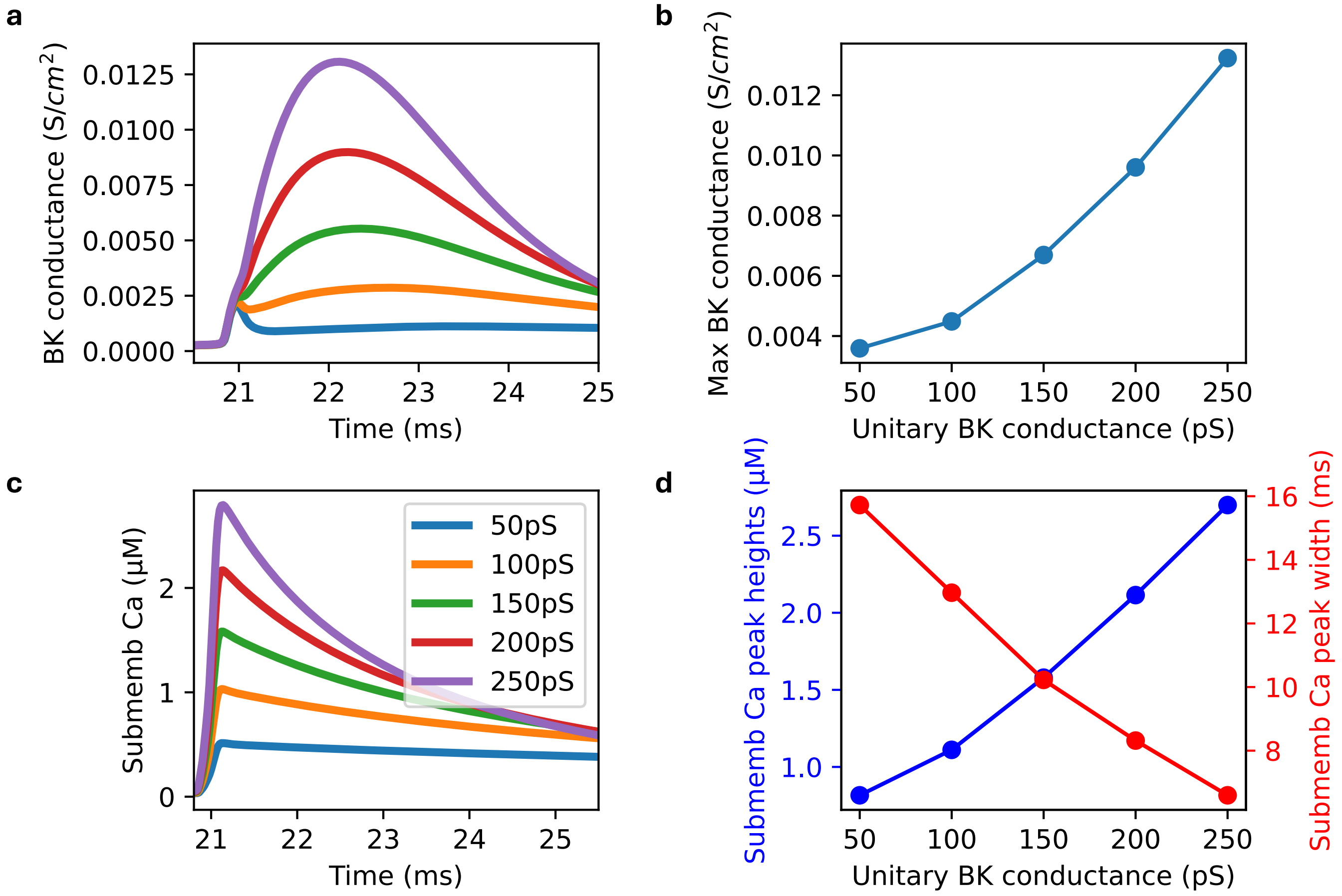


### *Supplementary Fig. 4: The calcium buffering effect of BK channels with this feature added to the NEURON model.*

*For one trial of the model shown in Fig. 5 where a calcium buffer is added to the model to mimic the calcium buffering effect of the BK channels, but without background synaptic noise:* ***a*** *The membrane BK conductance for the first, aligned spike.* ***b*** *The maximum membrane BK conductance over the full 500ms run of the model for different BK unitary conductances.* ***c*** *The calcium concentration in the submembrane shell on the first, aligned spike for the different BK unitary conductances tested.* ***d*** *The submembrane calcium maximum heights (blue) and widths (red) over the whole 500ms simulation for the different BK unitary conductances.*
